# A transgenic mutant mouse line accompanied by the complete deletion of interleukin-33 showed insulin and leptin resistances

**DOI:** 10.1101/416529

**Authors:** Taku Watanabe, Tomonori Takeuchi, Naoto Kubota, Tasuku Wainai, Keisuke Kataoka, Toshitaka Nakaya, Ayako Sugimoto, Takahiro Sato, Hiroshi Ohira, Ichizo Tsujino, Katsuyoshi Kumagai, Tetsuya Kubota, Chiaki Hasegawa, Kumpei Tokuyama, Kohjiro Ueki, Toshimasa Yamauchi, Masayoshi Mishina, Takashi Kadowaki

## Abstract

Interleukin (IL) −33 has been identified as a member of the IL-1 family. Members of the IL-1 family have been reported to be involved in the regulation of energy homeostasis and glucose metabolism. Homozygous transgenic mutant mice of FLP14 line, that we previously generated, unexpectedly developed mature-onset obesity and diabetes. Through genetic investigations, we found that insertion of the transgenes had resulted in complete deletion of the *Il33* gene. These obese male homozygous mutant mice exhibited hyperphagia with hyperleptinemia and insulin resistance caused by increased hepatic gluconeogenesis and decreased glucose uptake in skeletal muscle. As a result of examining preobese male homozygous mutant mice to investigate with the exclusion of the effect of obesity, hyperphagia with hyperleptinemina and insulin resistance caused by decreased glucose uptake in skeletal muscle were already observed, but the increased hepatic glucose production was not. To investigate whether the insulin resistance was caused by deletion of the *Il33* gene, we treated these preobese homozygous mutant mice with recombinant IL-33 protein and noted a significant improvement in insulin resistance. Thus, insulin resistance in these homozygous mutant mice was caused, at least in part, by IL-33 deficiency, suggesting a favorable role of IL-33 for glucose metabolism in the skeletal muscle.

---

The random insertion of transgenic fragments into the mouse genome by pronuclear microinjection is potentially mutagenic, and transgene-induced mutations were found in 5%–10% of transgenic mice^1-3^. There are examples of the accidental insertion of a transgene into a crucial genomic locus that yielded important information^4,5^. The present study began with the unexpected observation of the obesity phenotype in homozygous transgenic mutant mice of FLP14 line harboring the Flp recombinase^6^. We characterized the transgene integration site in this mutant mouse and found that the insertion of the transgenes was accompanied by a 124.2-kb deletion at the left end of the transgene and a 38.2-kb inversion and a 4.6-kb deletion at the right end. These mutations included the complete deletion of the *Interleukin* (*Il*) *33* gene.

Interleukin (IL) −33 (or IL1F11) has been identified as a functional ligand for the orphan receptor ST2, a member of the IL-1 receptor family^7^. Cytokines of the IL-1 family play an important role in immune regulation and inflammatory processes by inducing the expression of many effector proteins. In addition, members of the IL-1 family, IL-1and IL-18 have been reported to play a favorable role in the regulation of body weight and glucose metabolism^8-10^. IL-33 also has been shown to be pro-inflammatory in conditions such as autoimmunity^11^ and allergy^12^ and to be protective in conditions such as obesity^13-15^, atherosclerosis ^16^, and cardiac hypertrophy and fibrosis^17^.

The metabolic effects of IL-33 in obesity and diabetes have been investigated in murine models^13^. Administration of recombinant IL-33 protein to genetically obese diabetic (*ob/ob*) mice led to reduced adiposity, reduced fasting glucose and improved glucose and insulin tolerance. Furthermore, mice lacking endogenous ST2 fed high-fat diet had increased body weight and fat mass and impaired insulin secretion and glucose homeostasis compared to wild-type controls fed high-fat diet. These protective metabolic effects have been explained by the mechanisms that IL-33 induced production of Th2 cytokines (IL-5, IL-10, IL-13) and reduced expression of adipogenic and metabolic genes, and that IL-33 also induced accumulation of Th2 cells in adipose tissue and polarization of adipose tissue macrophages toward an M2 alternatively activated phenotype (CD206^+^). In addition, IL-33 reverses an obesity-induced deficit in visceral adipose tissue ST2^+^ T regulatory cells and ameliorates adipose tissue inflammation and insulin resistance^14^. In human, it has been reported that the serum levels of IL-33 was negatively associated with the body mass index and body weight in lean and overweight subjects^15^.

In the present study, we examined the glucose homeostasis and genetic profile of the homozygous mutant mice of FLP14 line that unexpectedly exhibited mature-onset obesity. Genetic analysis demonstrated complete deletion of the *Il33* gene in these mice, which raised the hypothesis that disappearance of IL-33 causes insulin and leptin resistance in this mutant mouse. To test this hypothesis, we investigated the impact of recombinant IL-33 protein supplementation on the glucose metabolism in the preobese male homozygous mutant mice.

## Results

### Mature-onset obesity in a homozygous transgenic mutant mouse of FLP14 line

Previously, we reported several transgenic mouse lines of the C57BL/6 strain carrying a Flp recombinase expression vector under the control of the human eukaryotic translation elongation factor 1*α* promoter^6^ (Fig. S1A). Among these lines, one homozygous mutant mice of FLP14 line (mutant) showed an obesity phenotype.

The body weights of the male mutant mice started to deviate from those of the male wild-type control (control) mice at 5 months of age and were increased by 13% at 6 months of age (Fig. 1A and B, left panels). On the other hand, the body weights of the female mutant mice started to deviate from those of the female control mice at 3 months of age and were markedly increased by 38% at 6 months of age (Fig. 1A and B, right panels). Gross appearance at sacrifice revealed that the fat mass in mutant mice was higher than that in the control mice (Fig. 1C). Moreover, marked fatty liver was observed in mutant mice (Fig. 1C). The weights of liver and visceral white adipose tissue (WAT) were also significantly increased in the mutant mice of both sexes compared with those in the control mice at 5 to 6 months of age (Fig. 1D).

**Figure 1.**
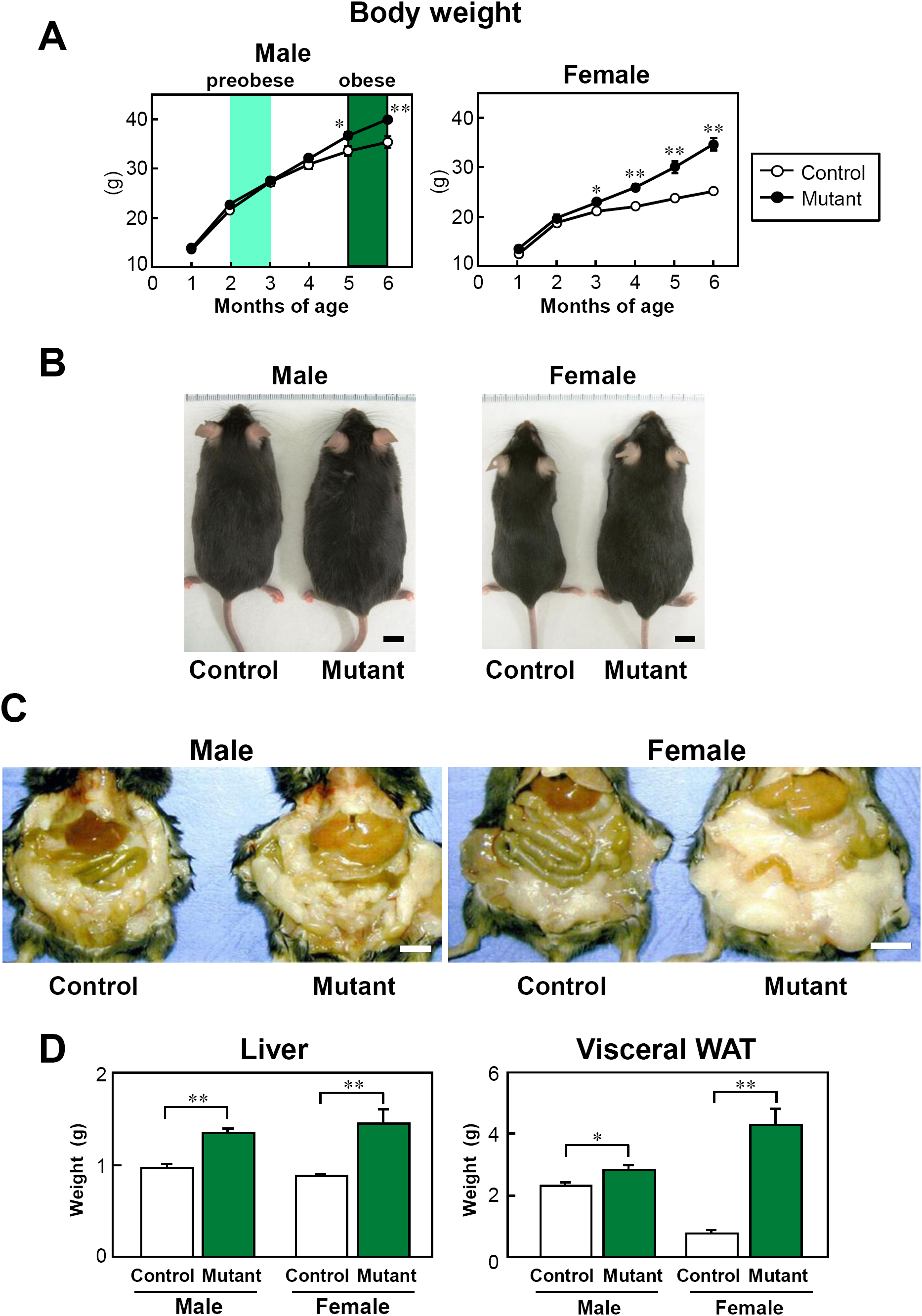
Mature-onset obesity in a homozygous transgenic mutant mice of FLP14 line. (**A**) Growth curves of the male (left) and female (right) homozygous transgenic mutant mice of FLP14 line (mutant) (filled circle) and wild-type control littermates (control) (open circle) fed on standard chow *ad libitum* (n = 8-11 per group). The obese phase was defined as 5 to 6 months of age, and the preobese phase was defined as 2 to 3 months of age. *P < 0.05, **P < 0.01. (**B**) Photographs showing the male (left) and female (right) control (left of each panel) and mutant (right of each panel) mice at 6 months of age. Scale bars, 1 cm. (**C**) Photographs showing the livers and intra-abdominal fat of the male (left) and female (right) control (left of each panel) and mutant (right of each panel) mice at 6 months of age. Scale bars, 1 cm. (**D**) Weight of the liver (left) and total visceral white adipose tissue (WAT) (right) of the control (white bars) and mutant (forest green bars) mice at 6 months of age. (n = 7-16 per group). *P < 0.05, **P < 0.01. Means ± SEM.

### The obese male mutant mice exhibited leptin and insulin resistances caused by both increased gluconeogenesis in liver and decreased glucose uptake in skeletal muscle

First, to examine the phenotype, the male mutant mice from 5 to 6 months of age were studied as the obese phase. The food intake of the male mutant mice was significantly increased compared with that of the control mice (Fig. 2A; left panel). The serum concentrations of leptin, which is an adipocytokine that regulates food intake, were significantly higher in the mutant mice than in the control mice (Fig. 2A; center panel). These association of hyperphagia and high leptin concentrations was suggestive of a leptin-resistant condition. Furthermore, the leptin/weight ratios in the mutant mice were also higher than those in the control mice (Fig. 2A; right panel). This result suggest that leptin resistance of the mutant mice was not a secondary induced by obesity induced but was primary. The serum cholesterol levels of the mutant mice were significantly higher than those of the control mice, although the serum triglyceride (TG) levels were indistinguishable (Fig. 2B; left and center panels). The serum adiponectin, which is an adipocytokine that regulates glucose and lipid metabolism, acting against diabetes and atherosclerosis, levels of the mutant mice were significantly lower than those of the control mice (Fig. 2B; right panel). The TG contents of both the muscle and the liver in mutant mice were significantly increased compared with those in control mice (Fig. 2C). The mutant mice exhibited elevated fasting blood glucose and serum insulin levels compared with those of control mice (Fig. 2D). These results were suggestive of an insulin-resistant condition. Therefore, we carried out hyperinsulinemic-euglycemic clamp studies to access the insulin resistance and to investigate whether the cause of the insulin resistance of the obese mutant mice is in the skeletal muscle and/or in the liver. In the hyperinsulinemic-euglycemic clamp study, the exogenous glucose infusion rates (GIRs) were significantly decreased in the mutant mice (Fig. 2E, left panel). Furthermore, the mutant mice exhibited not only a significant increase in the endogenous glucose productions (EGPs) but also a significant decrease in the rate of glucose disappearance (R_*d*_) (Fig. 2E, center and right panels). These results suggest that the obese mutant mice had insulin resistance caused by both increased hepatic glucose productions and decreased glucose uptakes by the skeletal muscle. These findings suggest that the genetic mutations of the mutant mice lead to obesity caused by hyperphagia, dyslipidemia, fatty liver and muscle, and diabetes with systemic insulin resistance.

**Figure 2.**
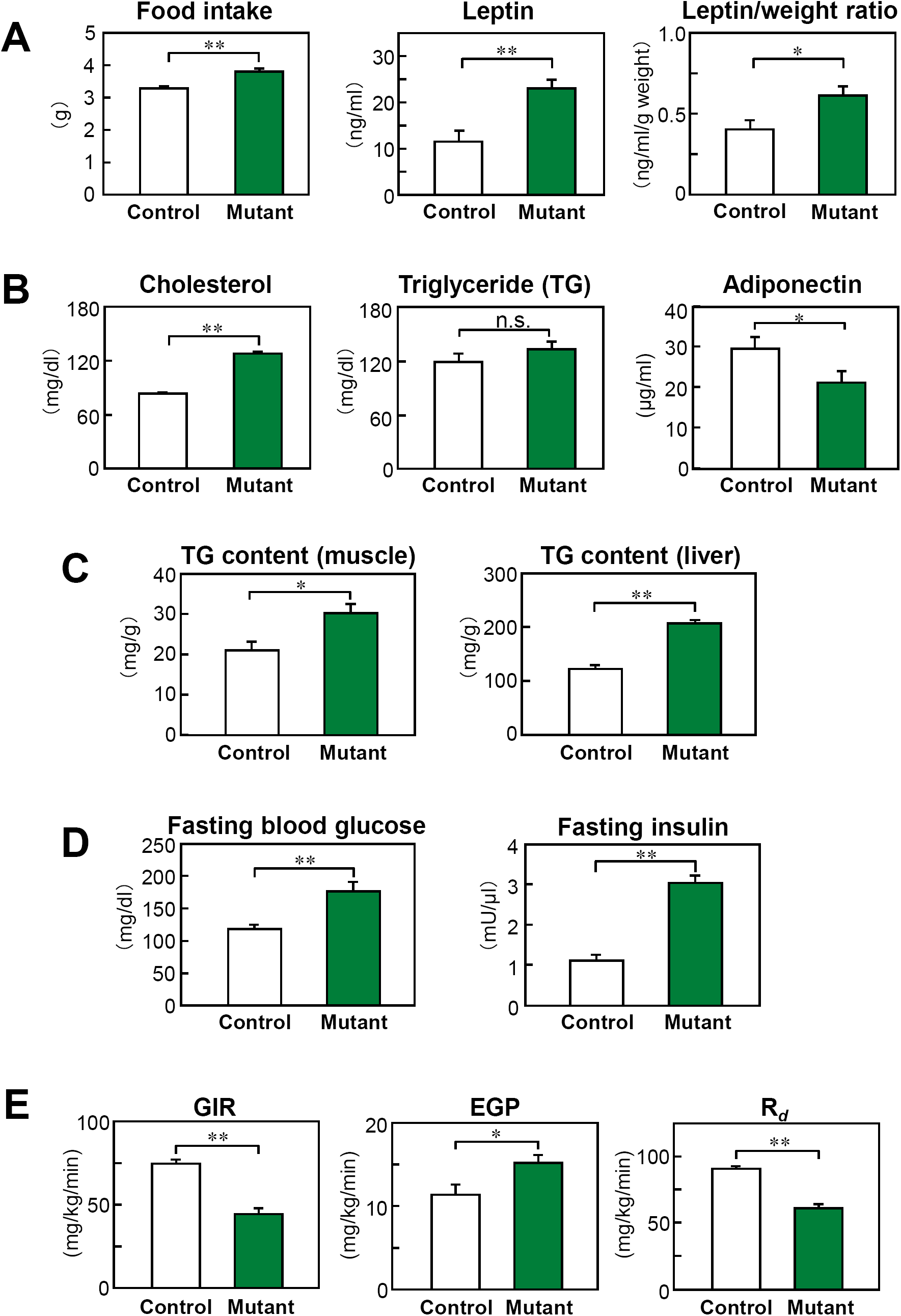
The obese male mutant mice from 5 to 6 months of age exhibited leptin resistance and insulin resistance caused by both increased gluconeogenesis in liver and decreased glucose uptake in skeletal muscle. (**A**) Food intake (left) (n = 7-12 per group), plasma concentrations of leptin (center) (n = 5-8 per group) and leptin/weight ratios (right) (n = 5-8 per group) of the control (white bars) and mutant (forest green bars) male mice at 6 months of age. *P < 0.05, **P < 0.01. (**B**) Plasma cholesterol (left, n = 5-6 per group), triglyceride (TG) (center, n = 6-8 per group) and adiponectin levels (right, n=5-7 per group) in the control and mutant mice. *P < 0.05; **P < 0.01; n.s., not significant. (**C**) Triglyceride (TG) content of the muscle (left, n = 5-10 per group) and of the liver (right, n = 5-8 per group) in the control and mutant mice. *P < 0.05, **P < 0.01. (**D**) Fasting blood glucose (left, n = 7-8 per group) and fasting insulin (right, n = 7-8 per group) levels in the control and mutant mice. **P < 0.01. (**E**) Glucose infusion rates (GIR) (left), endogenous glucose production (EGP) (center), and rates of glucose disappearance (R_*d*_) (right) of the control and mutant mice in the hyperinsulinemic-euglycemic clamp studies (n = 7-9 per group). *P < 0.05, **P < 0.01. Means ± SEM.

### The preobese male mutant mice already exhibited leptin and insulin resistances in skeletal muscle, but not in liver

Next, to exclude the effect of obesity, the young ‘preobese’ male mutant mice from 2 to 3 months of age who were the same weight as the control mice were studied. The food intake of the mutant mice was significantly increased compared with that of the control mice (Fig. 3A; left panel). The serum concentrations of leptin were significantly higher in the mutant mice than those in the control mice (Fig. 3A; right panel). To evaluate the energy expenditure, we investigated the oxygen consumption and found that it was indistinguishable between the mutant and control mice (Fig. 3B). These findings suggest that the preobese mutant mice already had leptin resistance that was independent of obesity, and that obesity was not due to reduced energy expenditure but due to hyperphagia caused by leptin resistance. The serum cholesterol and TG levels of the preobese mutant mice were significantly higher than those of the control mice (Fig. 3C; left and center panels). The serum adiponectin levels of the preobese mutant mice were indistinguishable from those of the control mice (Fig. 3C; right panel). The TG contents of both the muscle and the liver in mutant mice were not increased compared with those in control mice (Fig. 3D; left and center panels), and the weight of WAT in each genotype was indistinguishable (Fig. 3D; right panel). These results and previous experiments of obese mutant mice suggest that the decrease of serum adiponectin levels and fatty muscle and liver observed in the obese mutant mice were induced by obesity. However, importantly, the preobese mutant mice already exhibited elevated fasting blood glucose and serum insulin levels compared with those of the control mice (Fig. 3E). These results were suggestive of an insulin-resistant condition in the preobese mutant mice. We then next performed hyperinsulinemic-euglycemic clamp studies in the preobese mutant and control mice to investigate the insulin resistance of muscle and liver. We found that the GIRs were significantly decreased in the preobese mutant mice as compared to the control mice (Fig. 3F, left panel). Although the EGPs were indistinguishable between the mutant mice and control mice (Fig. 3F, center panel), the R_*d*_ was significantly decreased in the mutant mice as compared to the control mice (Fig. 3F, right panel). These results suggest that preobese mutant mice had insulin resistance caused by decreased glucose uptakes by the skeletal muscle, but the increased hepatic glucose production has not observed yet. To assess insulin signaling in the muscle of the preobese mutant mice, we conducted western blot analyses. Insulin-stimulated tyrosine phosphorylations of insulin receptor substrate-1(IRS-1) and insulin-stimulated phosphorylations of Akt were reduced in the muscle of preobese mutant mice as compared to that of control mice (Fig. 3G). These findings suggest that the genetic mutations in the mutant mice led to insulin resistance in skeletal muscle, but not in liver, and that this was independent of obesity.

**Figure 3.**
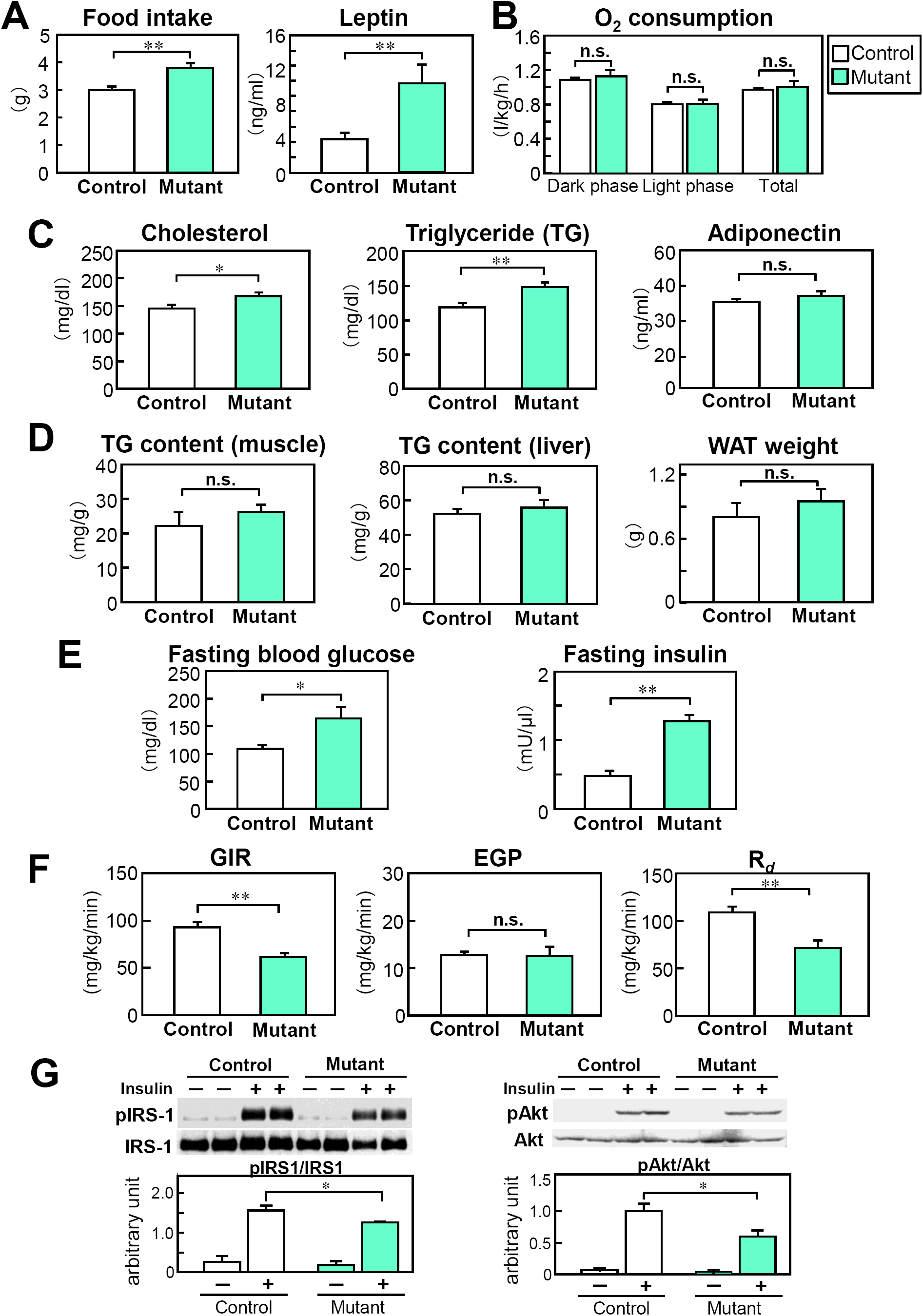
The preobese male mutant mice from 2 to 3 months of age already exhibited leptin resistance and insulin resistance caused by decreased glucose uptake in skeletal muscle. (**A**) Food intake (left, n = 8-12 per group), plasma concentrations of leptin (right, n = 12-13 per group) of the control (white bars) and mutant (mint green bars) male mice at 3 months of age. **P < 0.01. (**B**) Oxygen (O_2_) consumption in the control and mutant mice (n = 6-7 per group). n.s., not significant. (**C**) Plasma cholesterol (left) (n = 11-13 per group), triglyceride (TG) (center, n=6-8 per group) and adiponectin levels (right, n = 6-9 per group) in the control and mutant mice. *P < 0.05; **P < 0.01; n.s., not significant. (**D**) Triglyceride content of the muscle (left, n = 6-7 per group) and of the liver (center, n=7-10 per group), and weight of white adipose tissue (WAT) (right, n = 7-10 per group) in the control and mutant mice. n.s., not significant. (**E**) Fasting blood glucose (left, n = 6-8 per group) and fasting insulin levels (right, n = 12-13 per group) in the control and mutant mice. *P < 0.05, **P < 0.01. (**F**) GIR (left), EGP (center), and R_*d*_ (right) of the control and mutant mice in the hyperinsulinemic-euglycemic clamp studies (n = 6-12 per group). **P < 0.01; n.s., not significant. (G) Insulin-stimulated tyrosine phosphorylations of IRS-1 (left) and Akt (right) in the muscle of the control and mutant mice after the injection of insulin (n = 4-5 per group). Results are representative of three independent experiments. *P < 0.05. Means ± SEM.

### The *Il33* gene is completely deleted in mutant mice and recombinant IL-33 protein restored insulin resistance in the muscle of mutant mice

We hypothesized that the phenotype of obesity, and insulin and leptin resistances exhibited in these homozygous mutant mice was caused by transgenic insertional mutagenesis, because homozygous mice of the other five independent transgenic lines were not obese. Cumulative genotyping of mutant heterozygote crosses revealed that the manner of transgene transmission was consistent with Mendelian inheritance, indicating that the recessive obese phenotype was caused by disruption of a single locus in the genome with transgene insertion.

We used Southern blot analysis to analyze the copy number and structure of the transgene insertion site of the mutant mice. In an analysis of mutant genomic DNA cleaved with *Kpn*I, which cut both ends of the transgene, the tg-3’ probe detected two transgene-specific bands: a stronger fragment (∼3.4 kb) corresponding to a transgene expression vector and an additional weaker fragment (∼4.0 kb) (Fig. S1B). A comparison of these hybridization intensities with those of standard transgene expression vectors suggested that approximately 3 copies of the transgene were integrated (Fig. S1B). In an analysis of mutant genomic DNA cleaved with *Bgl*II, which cut the transgene at a single point, the tg-5’ probe or tg-3’ probe detected two transgene-specific bands in both cases: a stronger expected fragment (∼3.4 kb) and an additional weaker fragment (∼9.5 kb and ∼4.0 kb, respectively) (Fig. S1B). These results suggest that multiple copies of the transgene were integrated in tandem at a single locus.

We next cloned both the left-and right-flanking genomic sequences into which the transgene had been inserted. By using the basic local alignment search tool (BLAST), we determined that both the left-and right-flanking genomic sequences mapped to the mouse chromosome 19qC1 region containing three genes in the order of *Il33, Trpd52l3* and *Uhrf2* from centromere to telomere (Fig. 4A). The left-flanking sequence lies 48.8 kb upstream from the *Il33* transcription start site (Fig.S2), and the right-flanking sequence lies 0.4 kb upstream from the second exon of *Uhrf2* (Fig. S3). These two flanking sequences are 162.3 kb apart in the wild-type allele, while they are in opposite orientations in the mutant allele (Fig. 4A). Further analyses of the mutant allele surrounding the transgene integration sites using Southern blot analysis and genome walking indicated that the transgene integration caused the deletion of a 124.2-kb DNA fragment including the entire *Il33* gene at the ‘left integration (LI)’ site (Fig. 4A). At the ‘right integration (RI)’ site, the inversion of a 38.2-kb DNA fragment included the entire *Trpd52l3* gene and exon 1 of *Uhrf2*, and the deletion of a 4.6-kb DNA fragment included exon 2 of *Uhrf2* (Fig. 4A, Fig. S4). PCR of genomic DNA from the mutant mice with primers bracketing transgene insertion sites or the ‘right breakpoint (RB)’ region confirmed a map of the transgene insertion site (Fig. 4B).

**Figure 4.**
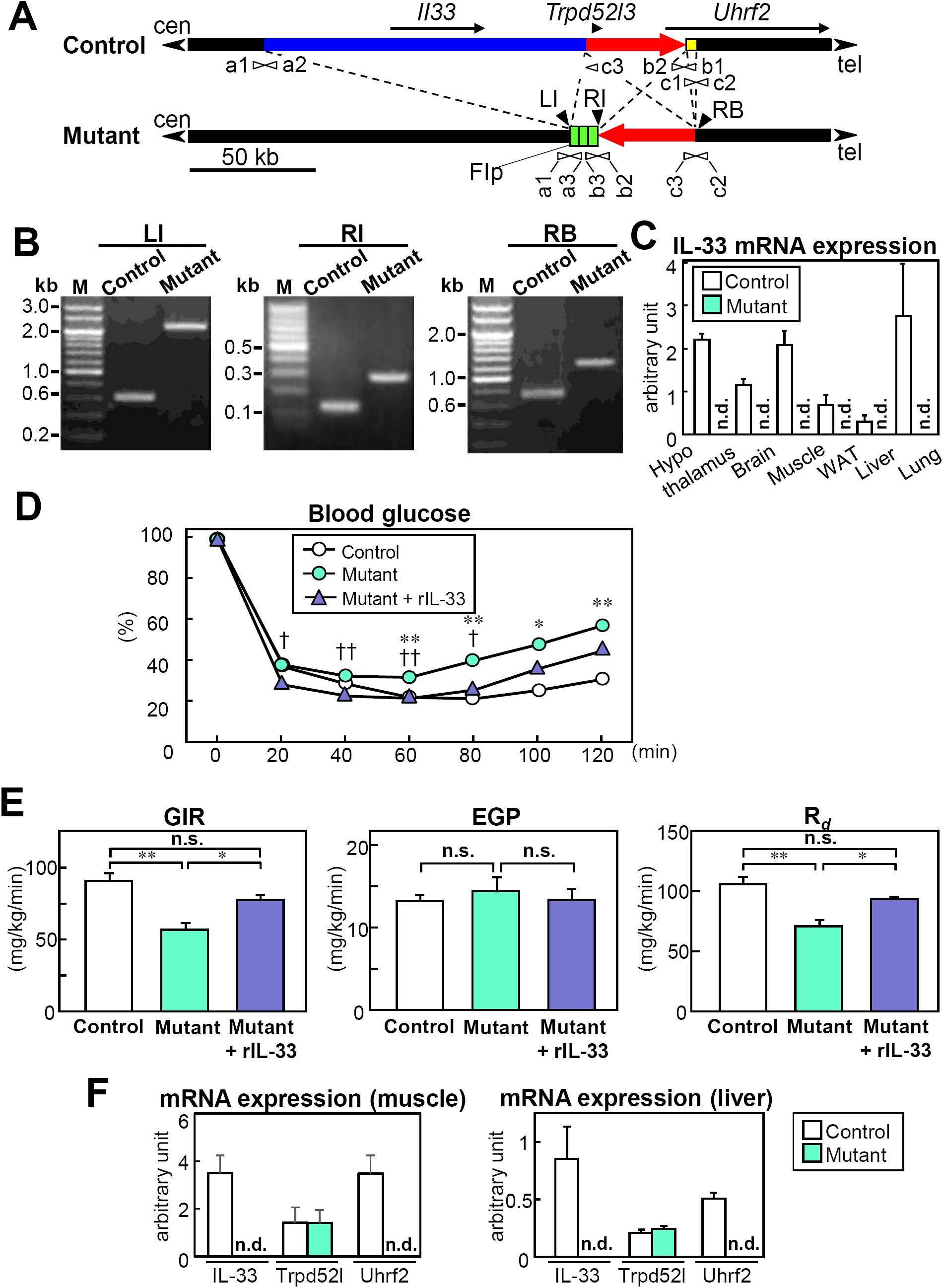
The *Il33* gene is completely deleted in the mutant mice and recombinant IL-33 protein restored insulin resistance in the muscle of the preobese mutant mice. (**A**) Schematic representation of the control and mutant allele surrounding the transgene integration site. Transgene integration caused the deletion of a 124.2-kb genomic DNA fragment (a blue region) at the left integration (LI) site of the mutant allele. It also caused the inversion of a 38.2-kb genomic DNA fragment with one break in the right breakpoint (RB) and the other break in the right transgene integration (RI) site (a red arrow) and the deletion of a 4.6-kb DNA fragment (a yellow region) at the right end of the mutant allele. The orientation of transcription is indicated by black arrows. White arrowheads indicate PCR primers (a1 and a3, b1-b3, and c1-c3) used for confirmation of the transgene insertion sites and breakpoint region. The orientation of the genomic DNA with respect to the centromere (cen) and telomere (tel) of chromosome 19 is indicated by black arrowheads. Flp, Flp recombinase expression vector pEF-NFLP. (**B**) PCR analysis of genomic DNA from the control and mutant mice using primers based on the inserted transgene and the flanking endogenous genomic DNA (left and center panels) or the flanking right breakpoint (right panel). Lane M contains size markers. (**C**) Real-time quantitative RT-PCR analysis in the various tissue of the control (white bars) and preobese mutant (mint green bars) mice (n = 7-10 per group). n.d., not detectable. (**D**) Insulin tolerance test (0.75 mU/g of body weight) results of the male control (white circles), the preobese mutant (mint green circles) and recombinant IL-33 (rIL-33) protein treated preobese mutant (light blue triangles) mice (n = 8-9 per group). The preobese mutant mice received an intraperitoneal injection of vehicle or rIL-33 at 1 hour before the insulin tolerance test. The control versus preobese mutant mice with vehicle: *P < 0.05, **P < 0.01; Vehicle versus rIL-33 treated preobese mutant mice: †P < 0.05, ††P < 0.01. (**E**), GIR (left), EGP (center), and R_*d*_ (right) of the male control (white bars), preobese mutant (mint green bars) and rIL-33 treated preobese mutant (light blue bars) mice in the hyperinsulinemic-euglycemic clamp studies (n = 6-12 per group). *P < 0.05; **P < 0.01; n.s., not significant. (**F**) Real-time quantitative RT-PCR analysis in the muscle (left) and the liver (right) of the control (white bars) and preobese mutant (mint green bars) mice (n = 7-10 per group). n.d., not detectable. Means ± SEM.

We identified 3 candidate genes: the *Il33* gene is completely deleted, the *Trpd52l3* gene is inverted, and the *Uhrf2* gene is inverted and partially deleted. Trpd52l3 is a mouse orthologue of the human tumor protein D52-like protein (TPD52L3 or hD55), which contains a coiled-coil motif and is strongly and uniquely expressed in human adult testis^18^. Uhrf2, which is also called Np95/ICBP90-like RING finger protein (NIRH) ubiquitin ligase, belongs to the ubiquitin PHD RING finger family and might behave as a tumor suppressor protein^19,20^. IL-33 is a member of the IL-1 cytokine family that induces production of T_H_2 cytokines^7^, and previous studies have indicated a role for the IL-1 family of cytokines in obesity^8-10^. Based on this information, we focused on *Il33* among the three candidate genes. We confirmed the deficiency of IL-33 mRNA expression in various tissues from mutant mice (Fig. 4C). As reported previously^7^, high levels of IL-33 mRNA expression could be found in the lung of the control mice.

We next treated the preobese mutant mice with recombinant IL-33 (rIL-33) protein before the insulin tolerance tests and the hyperinsulinemic-euglycemic clamp studies. Although the glucose-lowering effect of insulin was significantly impaired in the preobese mutant mice as compared to the control mice in insulin tolerance tests, the glucose-lowering effect of insulin was significantly restored by intraperitoneal administration of 2 µg of rIL-33 to the mutant mice 1 h before the studies (Fig. 4D). Similar to this result, in the hyperinsulinemic-euglycemic clamp studies, the GIRs were significantly restored by intravenous administration of 2 µg of rIL-33 to the preobese mutant mice 1 h before the studies (Fig. 4E; left panel). Although the EGPs were indistinguishable between the treated and untreated mutant mice (Fig. 4E, center panel), the R_*d*_ was significantly increased in the treated mutant mice as compared to the untreated mutant mice (Fig. 4E, right panel). These findings suggest that insulin resistance in preobese mutant mice was caused by IL-33 deficiency. However, by real-time quantitative RT-PCR analysis in the muscle and liver of the mutant mice as further examination about mutation, we were unable to detect the expression of IL-33 and Uhrf2 mRNA, though the expression of Trpd52l3 mRNA was comparable to the expression in those of control mice (Fig. 4F). Therefore, the possibility that the phenotype of mutant mice is affected by deletion of *Uhrf2* cannot be denied.

## Discussion

Here we showed that homozygous transgenic mutant mice of FLP14 line unexpectedly exhibit abnormalities characteristic of metabolic syndrome: obesity, hyperphagia, lipid abnormalities and insulin resistance. In these mutant mice, the insertion of the transgenes was accompanied by the complete deletion of the *Il33* gene. A hyperinsulinemic-euglycemic clamp study showed that the insulin resistance of obese male mutant mice was caused by increased hepatic gluconeogenesis and decreased glucose uptake in skeletal muscle. In contrast, preobese male mutant mice also exhibited insulin resistance and decreased glucose uptake in skeletal muscle; but, unlike obese mutant mice, they did not show increased hepatic glucose production. Administration of rIL-33 to the preobese male mutant mice improved insulin resistance and restored glucose uptake in the skeletal muscle, suggesting that insulin resistance in the mutant mice is caused by a deficiency of IL-33 and that IL-33 may play a pivotal role in glucose metabolism.

We identified two other genetic mutations (Fig. 4A) besides the complete deletion of the *IL33* gene in the mutant mice. One mutation is an inversion of *Trpd52l3* gene locus. Trpd52l3 is mouse orthologue of the human tumor protein D52-like protein, which is strongly expressed in adult human testis^18^. However, it notes that the expression of Trpd52l3 mRNA in the mutant mice was comparable to the expression in those of the control mice (Fig. 4F). Another mutation is an inversion and a partially deletion of *Uhrf2* gene locus, resulting in no expression of Uhrf2 mRNA in the mutant mice. Uhrf2 is a member of the ubiquitin PHD RING finger family and might act as a tumor suppressor protein^19^. Uhrf2 occupies a central position in the cell cycle network, has the capacity to ubiquitinate both cyclins D1 and E1, and connects the cell cycle machinery, the ubiquitin-proteasome system, and the epigenetic system^20^. To the best of our knowledge, however, no previous reports have indicated a relationship between these two genes and obesity or insulin resistance. Also, the almost complete improvement of insulin resistance after rIL-33 treatment suggests that these two genetic modifications have little impact on glucose metabolism.

The mutant mice in the present study exhibited insulin resistance. In general, the two major mechanisms of insulin resistance are increased hepatic glucose production and decreased glucose uptake in skeletal muscle. The former mechanism appeared to be caused by obesity itself rather than gene mutations in the mutant mice because glucose production in the liver was increased in the obese mutant mice (Fig.2E) but not in the preobese mutant mice (Fig. 3F). Also, serum adiponectin levels were significantly reduced only in the obese mutant mice as compared with the control mice (Fig. 2B, right panel). Adiponectin directly sensitizes the body to insulin^21,22^, and a reduced serum level of adiponectin might have prompted insulin resistance in the obese mutant mice. Likewise, increased hepatic TG content, which might have also caused insulin resistance^23-26^, was observed only in the obese mutant mice (Fig. 2C, left panel; Fig. 3D, left panel). On the other hand, decreased glucose uptake in the skeletal muscle appeared to be caused by genetic mutations independent of obesity because it occurred even in the preobese mutant mice (Fig. 3F). This insulin resistance in the skeletal muscle was also confirmed at the molecular level (Fig.3G). Furthermore, rIL-33 protein restored the decreased glucose uptake of skeletal muscle in preobese mutant mice (Fig. 4E, right panel), suggesting that the lack of IL-33 in the skeletal muscle was the primary cause of insulin resistance in the preobese mutant mice and, at least partly, in the obese mutant mice. In addition, in the control mice, the IL-33 expression level in the skeletal muscle was greater than in the liver (Fig. 4C). This result might explain why IL-33 deletion in preobese mutant mice affects glucose metabolism in the skeletal muscle, but not in the liver, in a manner that is independent of obesity.

The mutant mice in the present study developed mature-onset obesity (Fig. 1A) like other types of mice that are genetically deficient in IL-1 family members; mice deficient in IL-1 receptor I^ref9^ or IL-18^ref10^. In addition, IL-33 receptor ST2-deficient mice fed a high-fat diet also develop obesity^13^. Our obese mutant mice also showed increased food intake and leptin levels, indicating leptin resistance^27^. Importantly, we found the leptin levels in relation to body weight were higher in the obese mutant mice as compared with the control mice, whereas there was no such difference in IL-18 knockout mice^10^. Furthermore, leptin resistance also occurred in preobese mutant mice, which had increased food intake and serum leptin levels (Fig. 3A, right panel). Also, in the preobese mutant mice, the energy expenditure was not increased irrespective of the increased leptin levels (Fig. 3B). These findings suggest that IL-33 plays a key role in the regulation of body weight and that this regulation is at least partly achieved through leptin resistance. These findings suggest that hyperphagia induced by leptin resistance leads to a gradual increase in body weight, as the preobese mutant mice exhibited hyperphagia, but no change of energy expenditure. The role of IL-33 on body weight showed in the present study is consistent with the report that IL-33 is negatively associated with the body mass index and body weight in lean and overweight subjects^15^.

The effects of long-term rIL-33 administration on obesity have been reported by Miller and colleagues^13^. They treated spontaneously obese *ob/ob* mice with rIL-33 for 3 weeks. Consistent with the present study, administration of rIL-33 to *ob/ob* mice ameliorated glucose intolerance and insulin resistance. This treatment, however, resulted in decreased epididymal WAT weight, but not decreased body weight. It is possible that the effects of rIL-33 on obesity might have been reduced by the absence of leptin in *ob/ob* mice because the present study suggests that leptin plays a key role in the effect of IL-33 on obesity.

IL-33 has various effects. For example, IL-33 may be evolutionally preserved for the host defense against infections^11,28,29^. IL-33 has been implicated in regulating Th2-type innate immune responses^7^. IL-33 can also suppress the atherosclerotic process in ApoE-deficient mice^16^. Furthermore, IL-33 can exacerbate allergies^30,31^ and inflammation in arthritis^32,33^. Importantly, obesity, insulin resistance and type 2 diabetes are closely associated with chronic inflammation characterized by abnormal cytokine production, increased acute-phase reactants and other mediators, and activation of inflammatory signaling pathways^34,35^. Therefore, the metabolic effects of IL-33 showed in the present study may be explained by the alteration of inflammatory conditions^13,14^. Further investigations of the role of IL-33 are needed to clarify this issue.

Limitations of the present study include the possible influence of the mutations of *Trpd5213* and *Uhrf2*. In particular, unlike *Trpd5213, Uhrf2* was not detected in the mutant mice, suggesting that the lack of this gene product might have affected the results of this study. Studies using recombinant Uhrf2 are needed to further clarify this possibility. Also, we could not explore the underlying mechanisms of how the acute administration of rIL-33 improved insulin resistance and muscular glucose uptake on a molecular level. The use of IL-33-deficient mice instead of these mutant mice is indispensable for further studies. Lastly, we noted hyperphagia, even in the preobese mutant mice, which might have been caused by leptin resistance and may have affected the results. However, the insulin resistance observed in the preobese mutant mice had almost completely recovered immediately after rIL-33 administration, suggesting that IL-33 deficiency itself, rather than any changes caused by hyperphagia, was the primary cause of insulin resistance in our mutant mouse model.

In conclusion, this study suggests that insulin and leptin resistances in the homozygous transgenic mutant mice of FLP14 line is caused by a deficiency of IL-33 and that IL-33 has the possibility of being assigned a favorable role in glucose metabolism in the skeletal muscle.

## Methods

### Subjects

The Flp recombinase transgenic mouse line was generated by microinjection of C57BL/6N fertilized eggs with the Flp recombinase expression vector pEF-NFLP (Fig. S1A), as previously described^6^. Flp recombinase transgenic mice are now available at the RIKEN Bio-Resource Center through the National Bio-Resource Project of MEXT, Japan (www2.brc.riken.jp/lab/animal/detail.php?reg_no=RBRC01251). The homozygous (mutant) and wild-type littermates (control) derived from intercrossing the heterozygous mutant mouse were used for subsequent studies. Genotyping of the mutant mouse line by the polymerase chain reaction (PCR) was performed with primers 5’-CCTAGGCTAGCCTCAAACCTACAA-3’ (b1), 5’-CTCCCCACTCTTGGGAAT TTAGGT-3’ (b2), and 5’-TGAAAGGAAGCTCCACTGTCACCC-3’ (b3). Mice were fed *ad libitum* with standard chow, CE-2 (CLEA Japan Inc., Tokyo, Japan); the composition of the chow was as follows: 25.6% (w/w) protein, 3.8% fiber, 6.9% ash, 50.5% carbohydrates, 4% fat, and 9.2% water. Mice were kept in standard animal cages under a 12-h light: 12-h dark cycle. All experiments were performed with male and female littermates that were 2 to 3 months old in the preobese phase and 6 months old in the obese phase. All animal procedures were approved by the Hokkaido University Animal Experiment Committee (Approval number: 11-0115) and performed according to the guidelines for animal experimentation of Hokkaido University and the Animal Care and the Use Committee of the Graduate School of Medicine, the University of Tokyo (Approval: 1721T062).

The visceral WAT (mesenteric, retroperitoneal and epididymal fat pads in male mice or parametrial fat pads in the female mice) and liver were harvested and weighted.

### Plasma insulin, lipid, leptin and adiponectin measurements

The mice were deprived access to food for 16 hour before the measurements. Plasma insulin was measured with an insulin immunoassay (Morinaga Institute of Biological Science, Inc., Yokohama, Japan). Serum cholesterol and triglyceride (TG) (Wako Pure Chemical Industries Ltd., Osaka, Japan) were assayed by enzymatic methods. Plasma leptin and adiponectin levels were determined with a mouse leptin ELISA kit (R&D Systems, Minneapolis, USA) and a mouse adiponectin ELISA kit (Otsuka Pharmaceutical Co., Ltd, Tokyo, Japan), respectively.

### TG contents of the muscle and liver

For determining the TG content in the muscle and liver, the tissue homogenates were extracted and analyzed as described previously^36^.

### Hyperinsulinemic-euglycemic clamp study

Clamp studies were performed as described previously^37^ with slight modifications in insulin dose. In these studies, a primed continuous infusion of insulin (Humulin R) was administered (15.0 mU/kg/min for obese mice and 10.0 mU/kg/min for preobese mice). The rate of glucose disappearance (*R*_*d*_) was calculated according to nonsteady-state equations, and endogenous glucose production (EGP) was calculated as the difference between the *R*_*d*_ and the exogenous glucose infusion rate (GIR)^37^.

### Analysis of O_2_ consumption

Oxygen consumption was measured by using an O_2_/CO_2_ metabolism measurement device (Model MK-5000; Muromachikikai, Tokyo, Japan) as described previously^37^.

### Mouse recombinant IL-33 protein treatment study

We injected 2.0 µg mouse rIL-33 (Adipogen, San Diego, California, USA) or saline as the control intraperitoneally at 1 hour before the insulin tolerance test. In the hyperinsulinemic-euglycemic clamp studies, 2.0 µg rIL-33 protein or saline as the control was administered through the catheters at 1 hour before the studies.

### Tissue sampling for insulin signaling pathway study and western blot analysis

Mice were anesthetized after a 16-hour fast, and 0.05 units of human insulin (Humulin R) was injected via the inferior vena cava. After 5 min, the skeletal muscles were removed and immediately frozen in liquid nitrogen. Western blot analysis was performed as described previously^37^.

### Analyses of DNA

Genomic DNA was digested with *Kpn*I, which cut both ends of the transgene, and hybridized with the 2.1-kb *Bgl*II-*Bam*HI fragments from pEF-NFLP (tg-3’ probe) as a probe (Fig. S1). To estimate the copy number of the transgene, the intensity of the hybridizing signals was compared with the intensities of varying amounts of the 3.3-kb *Not*I fragment from pEF-NFLP as a standard.

To determine whether the transgenes were integrated at a single locus, genomic DNA digested with *Bgl*II, which cut the transgene at a single point, was hybridized with the 0.7-kb *Sal*I-*Bgl*II fragment from pEF-NFLP (tg-5’ probe) and tg-3’ probe.

A Genome Walker Universal Kit (Clontech, California, USA) was used to identify the mouse genomic sequences adjacent to transgene integration sites and a breakpoint^38^. To identify the left-flanking sequence of the transgene, genomic DNA from the mutant mouse was digested with *Hinc*II and ligated to adaptors supplied by the manufacturer. Nested PCR amplification of the fragment that contains flanking sequences of the left integration (LI) site was conducted by using the universal adaptor primers AP1 and AP2 and the following transgene-specific primers: 5’-CGTACTGCAGCCAGGGGCGTGGAAG-3’ and 5’-CTCTAGGCACCGGTTCAATTGCCGA-3’. Nested PCR-amplified products were cloned into pCRII-TOPO (Invitrogen, California, USA) and sequenced. PCR primers a1 (5’-AGACTAGAGCAGTAATGCTGGTAAG-3’), a2 (5’-TTTAGCCTACCGTTTTGTTCGTTAG-3’) and a3 (5’-CTCTAGGCACCGGTTCAATTGCCG A-3’) were used to confirm the LI site (Fig.4A).

Nested PCR amplification of the fragments that contain flanking sequences of the right integration (RI) site and surrounding sequences of the right breakpoint (RB) were similarly performed by using genomic DNA digested with *Ssp*I (for RI) or *Bgl*II (for RB) and the following gene-specific primers: RI, 5’-GAGCAGACAGGAGGAATCATGTCAG-3’ and 5’-TGAAAGGAAGCTCCACTGTCACCCT-3’; RB, 5’-TAAGGAAACACAAGGGCCATGGTTTTG-3’ and 5’-ACCTTCCAAGCTTCCATTGGCCAAATC-3’. The RI site was confirmed with PCR by using primers b1, b2 and b3. The RB region also was confirmed with PCR by using primers c1 (5’-AGGTTGTAGTTAGAGGCTGTGCACCACC-3’), c2 (5’-ACCTTCCAAGCTTCCATTGGCCAAATC-3’) and c3 (5’-CACAATCAGTTCCAAGACACAGCTACA-3’).

### Analyses of RNA

Total RNA was extracted from various tissues *in vivo*, and cDNA was synthesized as described previously^36^. mRNA levels were quantitatively analyzed by fluorescence-based reverse transcriptase-PCR. The reverse transcription mixture was amplified with specific primers by using an ABI Prism 7300 sequence detector (Applied Biosystems, California, USA). The primers used for *Il33, Trpd52l3, Uhrf2* and *cyclophilin* were purchased from Applied Biosystems. The relative expression levels were compared by normalization to the expression levels of *cyclophilin*.

### Statistical analyses

Statistical analyses were performed using JMP^®^Pro for Windows, version 10.0 (SAS Institute Inc., Cary, North Carolina, USA). Data were expressed as mean ± SEM. Differences between two groups were assessed by using unpaired t-tests. Data involving more than two groups were assessed by one-way ANOVA before Tukey-Kramer’s post-hoc test as appropriate to correct for multiple comparisons. All statistical tests were two-tailed. Statistical significance was set at *P*<0.05.

### Data availability

The dataset generated and analyzed in the current study is available from the corresponding author on reasonable request.

## Acknowledgements

We thank Haruka Nakahara, Chie Oda, Takanori Nomura, Eiko Kato, Mika Furuya, Yumiko Nakajima, Ken Osaki, Ritsuko Hoshino, Sayaka Sasamoto, Kayo Nishitani for their technical assistance, animal care, preliminary experiments and the preparation of this manuscript.

This work was supported by research grants from the Ministry of Education, Culture, Sports, Science, and Technology of Japan and the Japan Science and Technology Agency (12210007 and 17024011) (to T.T. and M.M.); Novo Nordisk Foundation Young Investigator Award 2017 (NNF17OC0026774), Aarhus Institute of Advanced Studies (AIAS)-EU FP7 Cofund programme (754513) and Lundbeckfonden (DANDRITE-R248-2016-2518) (to T.T.); a Grant-in-Aid for Scientific Research in Priority Areas ‘C’ (19591037) and ‘B’ (21390279) from the Ministry of Education, Culture, Sports, Science, and Technology of Japan (to N.K.); and a grant for Translational Systems Biology and Medicine Initiative (TSBMI) from the Ministry of Education, Culture, Sports, Science and Technology of Japan (to T.Ka); a Grant-in-Aid for Scientific Research in Priority Areas ‘A’ (16209030 and 18209033), and ‘S’ (20229008) from the Ministry of Education, Culture, Sports, Science, and Technology of Japan (to T.Ka.).

## Author Contributions

T.Wat. and T.T. designed and performed the experiments, analyzed the data, contributed to discussion and wrote the manuscript. N.K. contributed to discussion and reviewed/edited the manuscript. T.Wai., K.Ka., K.Ku., T.Ku., C.H. and K.T. performed the experiments. T.N., A.S., T.S. and H.O. contributed to discussion. I.T. contributed to discussion and reviewed/edited the manuscript. K.U., M.M. and T.Ka contributed to discussion and reviewed/edited the manuscript.

## Competing Interests

No potential conflicts of interest relevant to this article were reported.

## Supplemental Figure Legends

**Figure S1.** Multiple copies of the transgene were integrated in tandem at a single locus in a transgenic mutant mouse of FLP14 line. (**A**) Structure of the Flp recombinase expression vector. The Flp recombinase expression vector pEF-NFLP consists of the 1.3-kb promoter region of the human eukaryotic translation elongation factor 1 alpha 1 (EEF1A1) gene, the 1.4-kb SV40 large T antigen nuclear localization signal-Flp recombinase fusion gene (NFLP, a gray box) and the 0.6-kb human granulocyte colony-stimulating factor gene polyadenylation signal sequence (pA). Filled boxes indicate the first exon and part of the second exon of EEF1A1. The probes for Southern blots (tg-5’ and tg-3’) are indicated by white boxes. B, *Bgl*II; K, *Kpn*I; N, *Not*I. (**B**) Southern blot analysis of genomic DNA from the homozygous mutant mice. Left, *Kpn*I-digested genomic DNA hybridized with tg-3’ probe; middle, *Bgl*II-digested DNA hybridized with tg-5’ probe; right, *Bgl*II-digested DNA hybridized with tg-3’ probe.

**Figure S2.** Sequence across the left integration (LI) site in the mutant allele. The direction of the arrow boxes indicates the direction of the sequence on the wild-type chromosome 19, from centromere to telomere. Numbers in the arrow boxes indicate the corresponding sequence on the wild-type chromosome 19 (Ch 19). The left-end flanking sequence was juxtaposed to sequences derived from nt 761 to nt 2450 of pBluescript, which lie outside of the 3.3-kb *Not*I fragment from pEF-NFLP used for injection, possibly resulting from contamination of the injected fragment with partially digested plasmid DNA. The magenta arrowhead indicates the location of the junction. Arrows indicate PCR primers (a1 and a3) used for confirmation of the LI site.

**Figure S3.** Sequence across the right integration (RI) site in the mutant allele. The direction of the arrow boxes indicates the direction of the sequence on wild-type chromosome 19, from centromere to telomere. Numbers in the arrow boxes indicate the corresponding sequence on the wild-type chromosome 19 (Ch 19). The magenta arrowhead indicates the location of the junction. Arrows indicate PCR primers (b2 and b3) used for confirmation of the RI site.

**Figure S4.** Sequence across the right breakpoint (RB) in the mutant allele. The direction of the arrow boxes indicates the direction of the sequence on wild-type chromosome 19, from centromere to telomere. Numbers in the arrow boxes indicate the corresponding sequence on the wild-type chromosome 19 (Ch 19). The magenta arrowhead indicates the location of the break point. Arrows indicate PCR primers (c2 and c3) used for confirmation of the RB region.

